# Rewired RNAi-Mediated Genome Surveillance in House Dust Mites

**DOI:** 10.1101/065797

**Authors:** MH Mondal, NA Ortolano, KE Scott, CB Taylor, PB Klimov, AS Flynt

**Affiliations:** University of Southern Mississippi, Hattiesburg, MS, USA; University of Michigan, Ann Arbor, MI, USA; Tyumen State University, Tyumen, Russia

## Abstract

House dust mites are common pests with an unusual evolutionary history, being descendants of a parasitic ancestor. Transition to parasitism is frequently accompanied by genome rearrangements, possibly to accommodate the genetic change needed to access new ecology. Transposable element (TE) activity is a source of genomic instability that can trigger large-scale genomic alterations. Eukaryotes have multiple transposon control mechanisms, one of which is RNA interference (RNAi). Investigation of the dust mite genome failed to identify a major RNAi pathway: the Piwi-associated RNA (piRNA) pathway, which has been replaced by a novel small-interfering RNAs (siRNAs)-like pathway. Co-opting of piRNA function by dust mite siRNAs is extensive, including establishment of TE control master loci that produce siRNAs. Interestingly, other members of the Acari have piRNAs indicating loss of this mechanism in dust mites is a recent event. Flux of RNAi-mediated control of TEs provides a mechanism for unusual arc of dust mite evolution.

---

House dust mites are ubiquitous inhabitants of human dwellings, and are the primary cause of indoor allergy (Arlian, 1991). Dust mites have an unusual evolutionary history, descending from a parasitic ancestor (Klimov and B, 2013). Parasite genomes are typically highly modified; possibly to accommodate genetic novelty needed to productively interact with a host (Brookfield, 2011; Poulin and Randhawa, 2015). The sequence of events leading to adoption of a parasitic lifestyle may require genomic crisis to yield the rewired parasite genome. Dust mites represent an extreme case potentially experiencing a second round of genomic change to reacquire a free-living ecology.

Transposable element (TE) activity is a major source of genome instability (Fedoroff, 2012; Hedges and Deininger, 2007). RNA interference (RNAi), a process that employs small RNAs associated with Argonaute/Piwi (Ago/Piwi) proteins, directs TE silencing in multicellular organisms (Buchon and Vaury, 2006). In many animals the Piwi-associated RNA (piRNA) pathway is the primary RNAi-based defense (Crichton et al., 2014; Senti et al., 2015). In arthropods and vertebrates piRNAs are 25-30 nt long, and unlike other small RNAs, such as microRNAs (miRNAs) and small-interfering RNAs (siRNAs), they are not excised from double-stranded RNA (dsRNA) precursors by the RNase III enzyme Dicer (Weick and Miska, 2014). piRNAs are generated in two collaborative pathways: primary processing and the “ping-pong” amplification cycle (Czech and Hannon, 2016). Primary piRNAs are derived from designated single-stranded transcripts, which initiate the ping-pong cycle where Piwi proteins collaborate to capture fragments of TEs and convert them to new piRNAs (Iwasaki et al., 2015; Siomi et al., 2011). Ago/Piwi proteins may possess “slicer” activity where transcripts base-paired with a small RNA are cut 10 nucleotides (nt) from the 5’ end of the small RNA (Huang et al., 2014). TE transcripts processed into ping-pong piRNAs are produced by slicing, and therefore exhibit 10 nt 5’ overlaps with cognate, antisense piRNAs (Iwasaki et al., 2015). piRNA cluster master loci that generate primary piRNAs are comprised of TE fragments, and serve as catalogs of restricted transcripts (Malone et al., 2009). Loss of master loci integrity compromises TE repression and causes sterility.

While piRNA regulation of TEs is common in animals, it has been lost in several nematodes (Sarkies et al., 2015). In these species alternative mechanisms restrict TE mobilization that involve Rdrp (RNA dependent RNA polymerase) and Dicer. Conversion of TE transcripts by Rdrp into dsRNA substrates of Dicer results in siRNA generation. Ago proteins then associate with nascent TE transcripts, recruiting chromatin modulators including DNA methyltransferase. This process, RNA-induced transcriptional silencing (RITS) is common in plants and fungi (Klenov and Gvozdev, 2005; Pikaard, 2006; Verdel et al., 2009). Outside nematodes clades, RITS has not been observed in vertebrates or other ecdysozoans-potentially due to absence of Rdrp (Tomoyasu et al., 2008). One possible exception is chelicerae arthropods, a lineage where dust mites belong, which possess Rdrp proteins. RNAi pathways in chelicerates appear complex as they have both Rdrp and Piwi class Argonaute proteins (Grbic et al., 2011; Kurscheid et al., 2009). Here we investigate the status of small RNA pathways in the dust mite to shed light on TE control mechanisms in this highly derived organism.

We obtained a genome sequence for the American house dust mite *Dermatophagoides farinae* using Illumina and PacBio platforms (Supp Methods). Ago/Piwi proteins were identified in the *D. farinae* genome using amino acid sequences of seven Ago and six Piwi proteins from *Tetranychus urticae-the* closest relative of *D. farinae* with a high quality annotated genome (Grbic et al., 2011). Eight confident Ago homologs were found and annotated with RNA-Seq data (Supp. Fig. 1A)(Chan et al., 2015). Ago proteins from *T. urticae, D. melanogaster, C. elegans, and Ascaris suum* were compared to *D. farinae* Agos using amino acid sequences of Paz, Mid, and Piwi domains. Our phylogenetic analysis recovered two Ago family members likely involved in miRNA and siRNA pathways (Fig. 1A)(Meister, 2013). The remainder belongs to a divergent clade specific to dust mites (Fig. 1A). Surprisingly, none of the Agos from *D. farinae* belong to the Piwi clade. We examined *D. farinae* Agos for the presence of slicer activity. The DEDH slicer motif, which is common in metazoan Ago and Piwi proteins, were found in the *D. farinae* Ago1(miRNA) and Ago2 (siRNA). The divergent Agos have an uncommon DEDD catalytic motif (Fig. 1B). Orthologs containing a DEDD motif could be found in scabies *(S. scabiei),* social spiders *(S. mimosarum)* and *C. elegans* emphasizing the unusual nature of this Ago clade (Rider et al., 2015; Sanggaard et al., 2014; Tolia and Joshua-Tor, 2007).

**Figure 1:**
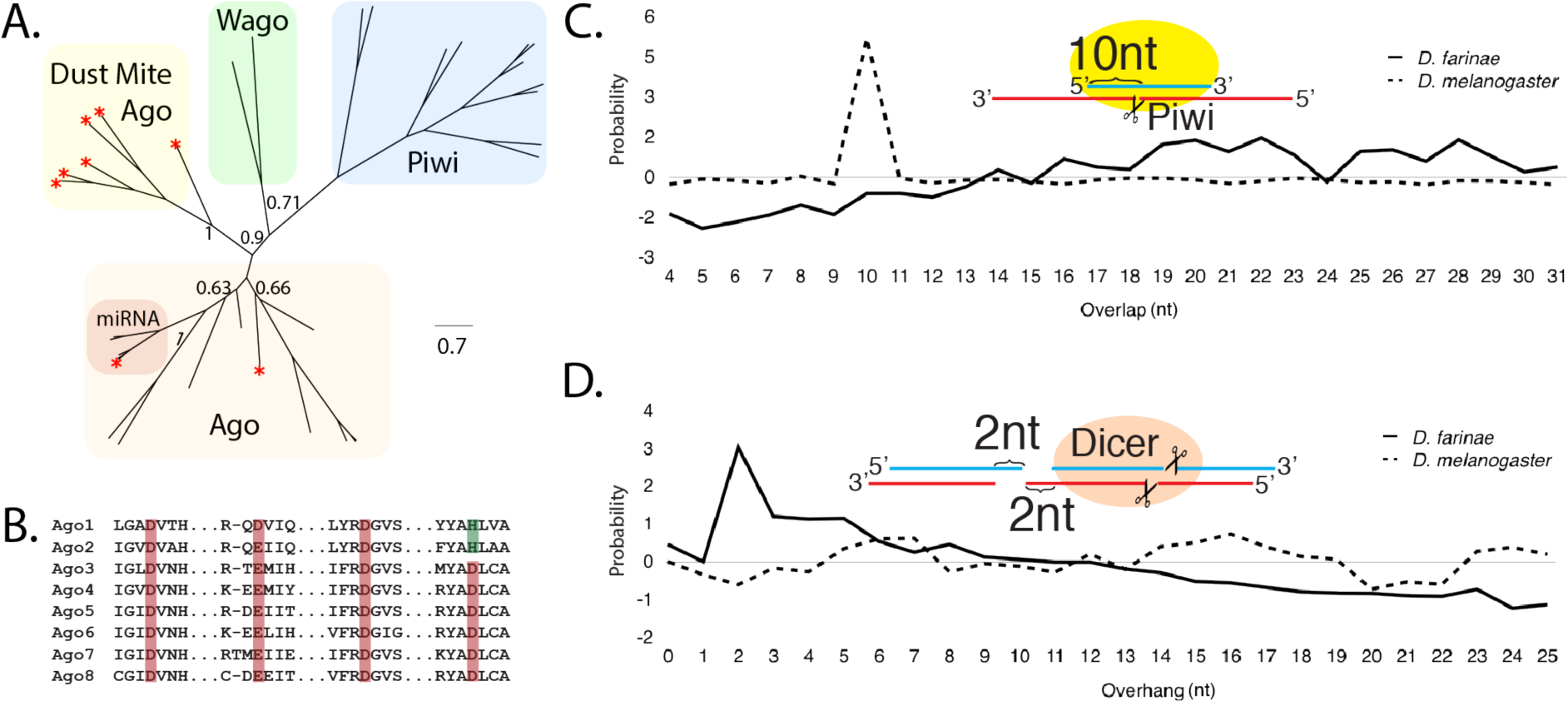
Absence of Piwi/piRNA pathway in dust mites. **A** Relationship of Ago/Piwi proteins from *D. farinae, Drosophila, C. elegans, and A. suum* using conserved Paz, Mid and Piwi domains. Asterisk indicates dust mite proteins. Only two Wago proteins included on tree for simplicity. Bootstrap values for major nodes indicated. **B.**Alignment of dust mite Ago “slicer” DEDH/D motif. Active site residues highlighted in red or green. **C.**Overlap probability for nucleotide overlaps for 24-31nt RNAs from *D. farinae* and *D. melanogaster.* **D.**Probability associated with overhangs of small RNA populations shown in C. Overhangs calculated by subtracting overlap length from read length for every size. Diagrams in C and D showing overlap and overhang characteristic of piRNA and siRNA biogenesis, respectively.

Next, a dust mite small RNAs library was analyzed for presence of piRNA-class transcripts. An algorithm that determines overlap probabilities was used to investigate whether the 10 nt overlap characteristic of ping-pong piRNAs was present in larger small RNAs (24-31nt) (Antoniewski, 2014). For fruit fly RNAs, we observed a peak at 10 nt overlap length, but not for dust mites (Fig. 1C). Investigation of overhangs revealed that 24-31 nt dust mite small RNAs possess a 2 nt overhang, a characteristic of Dicer cleavage (Fig. 1D). This indicates 24-31 nt small RNAs in dust mite are siRNAs produced by Dicer, and not piRNAs (Wen et al., 2014).

To further investigate the biology of dust mite siRNAs, we identified small RNA producing loci having >1000 read density that are longer than 200 nt. We found ~400 loci that produced 75% of the small RNAs mapping to the *D. farinae* genome (Supp. Table 1&2). The identities of the regions were determined using blast2go (Conesa et al., 2005). Nearly a third of the loci were rRNA or mRNA. The remainder showed homology to TEs, along with a large cohort of loci that lacked similarity to known sequences. These unknown elements produce over half of the reads mapping to longer loci. The remaining 21% of small RNAs align to TEs. To determine whether any of the TE loci might be processed into piRNAs we investigated if strand bias was present. Inspection of loci revealed reads emanating from both strands when considering all mapping events or only unique hits (Supp. Table 2). This Indicates that like the ping-pong pathway, primary piRNAs are also absent from dust mites.

To investigate the role of dust mite small RNAs in genome surveillance we sought to characterize biogenesis of small RNAs across all TEs. Nearly 700 TE loci were annotated using RepeatMasker (Smith et al., 2007). Compared to the bimodal distribution of TE small RNAs in *Drosophila* we found dust mite small RNAs that mapped to TE loci were unimodal with a small portion in the 26-31 nt size range (Supp. Fig. 3). The dust mite small RNAs also did not exhibit a strong 1U bias typical for piRNAs in *Drosophila* (Fig. 2A). Instead we observe an equal distribution of A and U at the first position. Furthermore, the TE derived small RNA show a 2 nt overhang bias (Fig. 2B). Together this suggests siRNAs are the main mode for controlling TEs in dust mites, accommodating the apparent loss of piRNAs.

**Figure 2:**
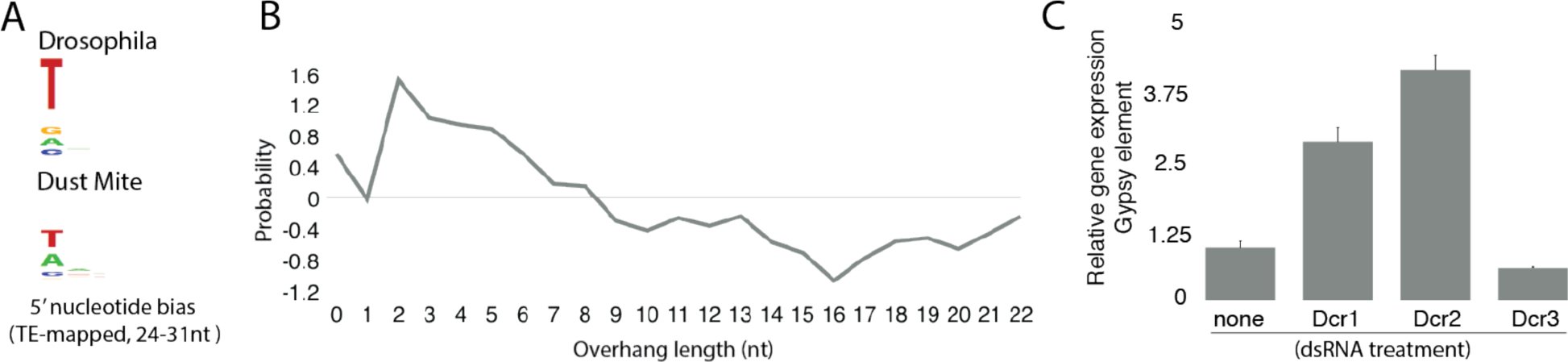
TE derived small RNAs are Dicer dependent. **A.** Seqlogo showing nucleotide bias at the 5’ end of 24-31nt small RNA that map to TEs in *Drosophila* and dust mites **B.**Probability of overhangs associated with small RNAs mapping to annotated TEs in the *D. farinae* genome. **C.**Expression of a gypsy element following dsRNA soaking against the three dust mite Dicers. Error bars represent SEM.

To verify whether TEs are controlled by siRNAs we sought to elicit RNAi against dust mite Dicer proteins by delivery of dsRNA trigger. Dust mites tolerate being soaked in dsRNA; after which, knockdown of target gene expression can be observed (Supp. Fig. 4). In the *D. farinae* genome we found three Dicers (Supp. Fig. 1B). Depletion by RNAi of Dcr1 and Dcr2 but not Dcr3 caused derepression of a Gypsy element, a known target of piRNAs in flies (Fig. 2C)(Brennecke et al., 2007). We identified moderate to strong nuclear localization signal (NLS) in Drc1 and Dcr2, but not in Dcr3 (Supp. Table 3). Several of the Ago proteins and an Rdrp homolog also possess NLS domains. This suggests that dust mites may be repressing TE activity through a nuclear RITS-type mechanism acting on nascent transcripts.

Investigation of RNAi in dust mites revealed loss of the piRNA pathway and replacement by siRNAs. This is similar to observations in nematodes where piRNAs were lost multiple times during this group’s radiation (Sarkies et al., 2015). The loss of piRNA activity in dust mites and nematodes is likely to be tolerated due to compensation by amplifying siRNAs produced by Rdrp (Sarkies et al., 2015). The collective function of dust mite Rdrps, however, appears to be distinct from nematodes, as only processive versions are present (Supp. Fig. 5). This suggests the *de novo* siRNA pathway is not presents in mites, which is consistent with our observation that small RNAs in dust mites are Dicer products (Figure 2). Substantial Rdrp activity does appear to be present in dust mites; dsRNA soaking causes elevation of target mRNA that accompanies decrease in protein accumulation (Supp. Fig. 4B).

Dust mites also differ from nematodes that lack piRNAs in the organization of siRNA producing loci. A key feature of piRNA biology is the cataloging of restricted sequences into master loci. In the nematode lineages lacking piRNAs master loci also appear to be absent (Sarkies et al., 2015). This is not the case in dust mites (Fig. 3A). Three loci were discovered that span 62 kb, contain sequences from multiple varieties of TE, and exhibit homology to 70% of TE mapped small RNAs. Similar regions could not be found in the S. *scabiei* genome (Rider et al., 2015). Poor conservation is a characteristic of piRNA master loci (Shi et al., 2013). The dust mite loci generate siRNAs, arising from an apparent dsRNA precursor as both strands of the loci show similar rates of read mapping (Fig 3A). It is unclear whether these regions are piRNA master loci that converted to siRNA production or are the result of genome rearrangements that occurred following the loss of the piRNA pathway function.

**Figure 3:**
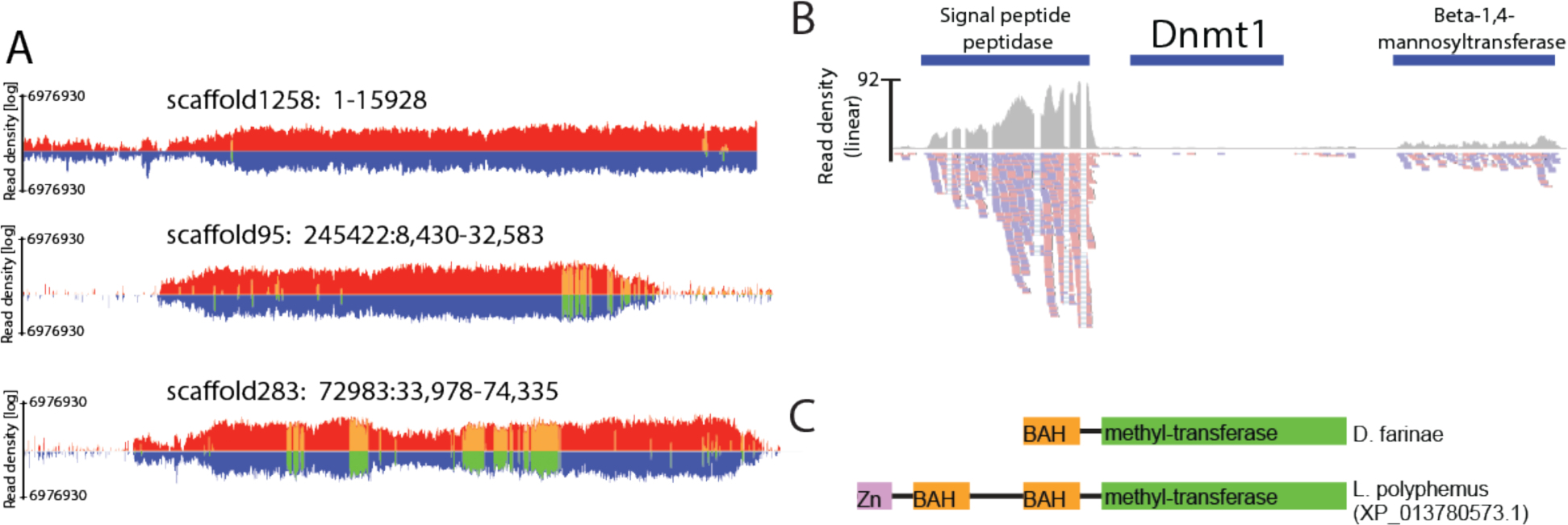
Characteristics of small RNA-mediated TE control in dust mites. **A.** siRNA producing TE-control master loci. Read density of all mapping events to the positive strand in red, negative strand blue. Density of uniquely mapping reads in yellow for positive strand and green for negative strand. **B.** Expression of dust mite DNA (cytosine-5)-methyltransferase 1 (Dnmt1) in mixed stage RNA-Seq data. Blue bar represents dust mite Dnmt1 locus in the scaffold. Read density in region shown as grey plot. Reads mapping below. Plus strand mapping in red, minus strand mapping in blue. **(C)** Domain structure of truncated **D.** farinae Dnmt1 and an intact ortholog from Limulus polyphemus.

Nematodes ultimately use siRNAs to drive DNA methylation (Sarkies et al., 2015). Dust mites further differ from nematodes in this regard. We found a single DNA methyltransferase in the *D. farinae* genome, a Dnmt1 homolog. It is likely a pseudogene as it appears to be truncated and shows little evidence of expression (Fig 3B,C). Furthermore, substantial methylation was not observed in dust mites using an immunoblotting method (Supp. Fig. 5). This is consistent with loss of DNA methylation of TEs in arthropods, and highlight the distinct, derived nature of small RNA-mediated genome surveillance in dust mites (Yan et al., 2015).

This work provides insight into the elaborate nature of RNAi in chelicerates. Most members of this clade appear to have both Piwi proteins and Rdrps (Grbic et al., 2011; Kurscheid et al., 2009; Sanggaard et al., 2014). However, there doesn’t appear to be much benefit to having both pathways as many chelicerate genomes are heavily colonized by TEs (Gulia-Nuss et al., 2016). The loss of the piRNA pathway in dust mites probably occurred in the parasitic ancestor. Inspection of the scabies genome failed to uncover piwi homologs; however, without transcriptome information systematic annotation of scabies Ago proteins was not possible (Rider et al., 2015). It is not clear whether loss of piRNAs accompanied transition to parasite ecology or occurred later in the group’s radiation.

Flux of small RNA-mediated TE surveillance correlates with evolutionary innovation; for example, higher arthropods lost Rdrp in favor of piRNA control of TE (Maida and Masutomi, 2011). This also occurred when vertebrates diverged from basal chordates (Putnam et al., 2008). In both cases loss of Rdrp accompanied innovation in body plan and sensory organs. In vertebrates whole genome duplication occurred twice following descent from an Rdrp expressing ancestor affirming a period of genome instability (Putnam et al., 2008). TE activity may be fortuitous for adaptation, and dramatic evolutionary changes require extreme events such as perturbation of surveillance mechanisms.

